# Deletion of an evolutionarily conserved TAD boundary compromises spermatogenesis in mice

**DOI:** 10.1101/2024.07.08.602428

**Authors:** Ana C. Lima, Mariam Okhovat, Alexandra M. Stendahl, Jake VanCampen, Kimberly A. Nevonen, Jarod Herrera, Weiyu Li, Lana Harshman, Ran Yang, Lev M. Fedorov, Katinka A. Vigh-Conrad, Nadav Ahituv, Donald F. Conrad, Lucia Carbone

**Author notes:** These authors contributed equally to the project (Ana C. Lima, Mariam Okhovat). Joint supervision (Donald F. Conrad, Lucia Carbone).

## Abstract

Spermatogenesis is a complex process that can be disrupted by genetic and epigenetic changes, potentially leading to male infertility. Recent research has rapidly increased the number of protein coding mutations causally linked to impaired spermatogenesis in humans and mice. However, the role of non-coding mutations remains largely unexplored. As a case study to evaluate the effects of non-coding mutations on spermatogenesis, we first identified an evolutionarily conserved topologically associated domain (TAD) boundary near two genes with important roles in mammalian testis function: *Dmrtb1* and *Lrp8*. We then used CRISPR-Cas9 to generate a mouse line where 26kb of the boundary was removed including a strong and evolutionarily conserved CTCF binding site. ChIP-seq and Hi-C experiments confirmed the removal of the CTCF site and a resulting increase in the DNA-DNA interactions across the domain boundary. Mutant mice displayed significant changes in testis gene expression, abnormal testis histology, a 35% drop in the estimated efficiency of spermatogenesis and a 28% decrease in daily sperm production compared to littermate controls. Despite these quantitative changes in testis function, mutant mice show no significant changes in fertility. This suggests that non-coding deletions affecting testis gene regulation may have smaller effects on fertility compared to coding mutations of the same genes. Our results demonstrate that disruption of a TAD boundary can have a negative impact on sperm production and highlight the importance of considering non-coding mutations in the analysis of patients with male infertility.

## INTRODUCTION

Spermatogenesis is a complex and meticulously regulated process essential for male fertility. During spermatogenesis, hundreds of millions of spermatozoa are generated daily from a renewable population of spermatogonial stem cells in the seminiferous tubules of the testis. In humans, infertility affects ∼15% of couples globally, and male factors are estimated to contribute to half of these cases^1^. Nearly half of male infertility cases remain unexplained after clinical examination, and much work is being done to understand molecular mechanisms underlying these unexplained cases. After nearly a decade of exome-sequencing studies, monogenic genetic causes can now be identified in about 20% of these “unexplained” cases^2-4^. Due to the nature of exome sequencing, most genetic causes identified in these cases are protein-coding. The first whole-genome sequencing (WGS) studies of male infertility are emerging^5,6^, but interpretation of non-coding variants remains a massive challenge for this field and human genetics in general. Rigorous characterization of non-coding mutations in mouse models are needed to better understand the impact of regulatory variation on reproductive traits, and, in turn, draw generalizations that can be applied to the interpretation of patient genomes.

In three-dimensional space, the genome is arranged into non-random structural units called Topologically Associated Domains (TADs)^7^. TAD boundaries are often bound by CTCF (CCCTC-binding factor) and cohesin complexes, and are crucial in maintaining appropriate spatial and temporal coordination of gene expression by preventing ectopic interactions across neighboring TADs. TAD boundaries play a fundamental role in developmental gene regulation, and disruption of their organization, often through deletion or misplacement, may lead to developmental disorders, such as structural birth defects^8,9^ and cancer^10,11^. The evolutionary conservation of a TAD boundary appears to be linked to the functional importance of nearby genes, with many highly conserved TAD boundaries flanking developmental genes, whose misregulation can be lethal in mouse models and may lead to genomic disorders in patients^12,13^.

A number of recent studies have mapped TADs in mouse testicular germ cells, and some general conclusions have emerged^14-16^; TADs can be identified in all phases of spermatogenesis examined so far. Spermatogonia are similar to somatic cells (*e*.*g*., fibroblasts) in the number and strength of their TAD boundaries^16^. The number, location, and strength of TADs changes through spermatogenesis, with the number of TADs dropping dramatically during meiotic prophase, increasing sharply in round spermatids, and stabilizing in spermatozoa. Like somatic cells, CTCF binding is enriched at TAD boundaries in germ cells.

Given the importance of TAD boundaries to developmental gene regulation, we conducted a case study to investigate the link between TAD organization and spermatogenesis. We identified and analyzed a TAD boundary that neighbors important testis-expressed genes, is conserved across human and four distant mammalian species (rhesus, rabbit, dog, and mouse), and overlaps synteny breakpoints between human and gibbon^12^, a small ape whose genome has undergone a high rate of evolutionary rearrangements^17^. Based on this evidence, this TAD boundary has likely been preserved for >80 million years and despite genomic rearrangement events, indicating it plays a critical role in regulating nearby genes. Located near this conserved TAD boundary are two genes involved in spermatogenesis, *Dmrtb1* and *Lrp8*. Doublesex- and mab-3-related transcription factor B1 (DMRTB1, synonym: DMRT6) is a member of the highly conserved DMRT family of transcription factors, which plays a crucial role in sex determination and gamete development across species^18^. Particularly, DMRTB1 seems to regulate spermatogonial differentiation and meiotic progression in mice^19^, and it has been associated with spermatogenic impairment in humans^20^. The LDL Receptor Related Protein 8 gene (*Lrp8*, synonym: *ApoER2*) encodes a key protein in testicular selenium metabolism, and its deficiency in mice leads to epididymal sperm abnormalities^21^. To investigate if evolutionary conservation of our candidate boundary reflects its involvement in spermatogenesis and fertility, we removed this boundary using CRISPR-Cas9 in a mouse model. Despite being fertile, the mice homozygous for the deletion showed significant changes in testicular gene expression, testis histology and daily sperm production following the deletion, demonstrating that disruption of evolutionary conserved TAD organization can negatively impact sperm production. Studying TAD boundaries in the context of sperm production not only deepens our understanding of fundamental biological processes, but also holds promise for improving genetic analysis of WGS from patients with unexplained male infertility.

## RESULTS

### Generation of a knock-out mouse model to investigate the importance of TAD organization in spermatogenesis

We selected our candidate TAD boundary from a set of 18 evolutionarily conserved TAD boundaries that were previously reported to overlap human vs. gibbon breakpoints of evolutionary chromosome rearrangements and be shared across genomes of six different mammalian species (human, gibbon, rhesus, rabbit, dog, and mouse)^12^. Our selected boundary is located between genes relevant to spermatogenesis, *Dmrtb1* and *Lrp8*, but does not overlap with any protein-coding genes, minimizing the risk that its deletion would disrupt integrity of any genes (Fig. 1A). To keep the CRISPR-Cas9 deletion size relatively small, we targeted a 23kb region encompassing an evolutionarily conserved CTCF binding site (Fig. 1A) within our candidate conserved TAD boundary (mm10, chr4:107716841-107739906). We validated the deletion in mutant mouse lines (*Dmrtb1_B*^*-/-*^) via PCR, Southern blot and Sanger sequencing (Supplementary Table 1, Supplementary Fig. S1). Bulk testis CTCF ChIP-seq also confirmed successful removal of the targeted CTCF binding site in *Dmrtb1_B*^*-/-*^ testis, without any other notable changes in nearby CTCF binding sites (Fig. 1B).

**Figure 1.**
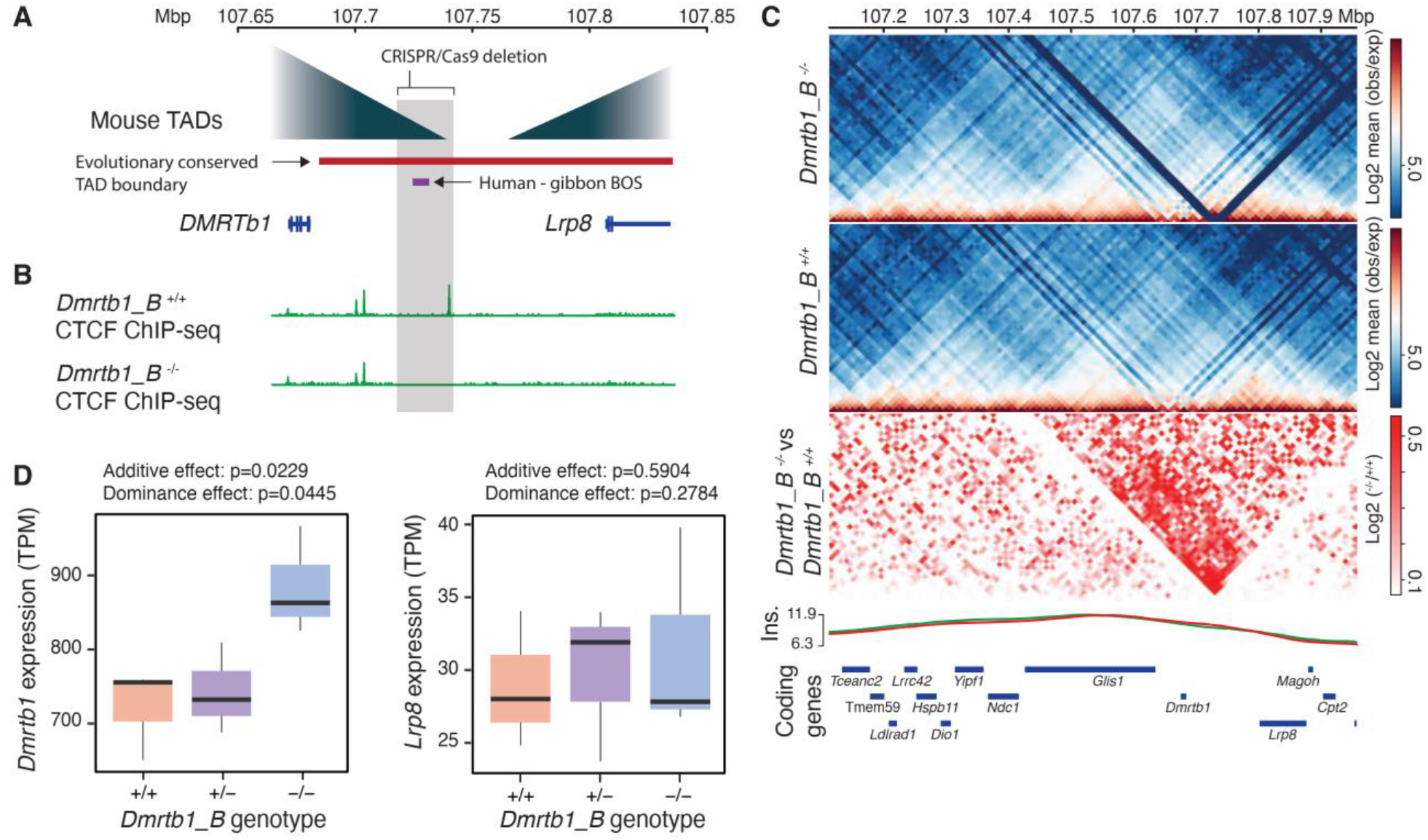
TAD boundary selection, followed by generation and validation of *Dmrtb1_B*^-/-^ mice. A) Schematic view of the conserved TAD boundary, the gibbon vs. human break of synteny (BOS), and CRISPR deletion in *Dmrtb1_B*^*-/-*^ mice. B) CTCF ChIP-seq fold-enrichment tracks show successful removal of the targeted CTCF peak in testis of *Dmrtb1_B*^*-/-*^ mice. C) Top two contact heatmaps show Capture Hi-C data from testis of *Dmrtb1_B*^-/-^ and *Dmrtb1_B*^*+/+*^ mice around the CRISPR deletion site. Bottom heatmap represents log difference between two heatmaps above and reveal mild increase in interaction across the deletion site, while insulation score curves below (green = *Dmrtb1_B*^*+/+*^ and red = *Dmrtb1_B*^*-/-*^) indicate no major change in genome conformation following the deletion. Positions of coding genes are shown below. D) Testicular *Dmrtb1* and *Lrp8* expression is shown for different genotypes of the *Dmrtb1_B* line based on RNA-seq data. TPM= transcript per million.

### Deletion of an evolutionary conserved TAD boundary near spermatogenesis genes results in gene expression changes in mouse testis

To evaluate the effects of our deletion on genome conformation, we performed capture Hi-C using a set of probes covering the region of interest and a control region on bulk testis (Methods). Although we did not identify major changes in TAD organization or insulation curves between homozygous mutant (*Dmrtb1_B*^*-/-*^) and wild-type mice (*Dmrtb1_B*^*x/+*^), we did detect a mild increase in chromatin interaction frequency across adjacent TADs, following deletion of the conserved boundary (Fig 1C). To determine the consequences of the deletion on gene expression, we performed RNA-seq on bulk testis of homozygous and heterozygous *Dmrtb1_B* mice (*DMRTB1_B*^*-/-*^ and *Dmrtb1_B*^*+/-*^), as well as wild-type controls (*DMRTB1_B*^*+/+*^). Genome-wide differential gene expression analysis did not reveal any significant results, likely due to the small sample sizes (*n*=3 per genotype). However, eQTL analysis on genes located within 1Mb of the deletion site (Fig. 1D) revealed a significant (p=0.023; Wald test) dose-dependent increase in *Dmrtb1* expression in *Dmrtb1_B*^*-/-*^ compared to *Dmrtb1_B*^*+/+*^ mice, consistent with the mild increase in chromatin interaction frequency detected in the Hi-C data (Fig. 1C). No other differentially expressed genes were detected, including *Lrp8*, the other gene adjacent to the deleted TAD boundary (Fig. 1D). Given that *Lrp8* has strong expression in Sertoli cells^22^, which represent a small fraction of the total testis cell population, and low expression in round spermatids^22^, we believe RNA-seq of bulk testis will have limited ability to detect subtle expression changes in this gene.

To validate these findings and determine if gene expression differences are specific to a given stage of spermatogenesis, we performed RNA *in situ* hybridization with RNAscope™ (Advanced Cell Diagnostics) using probes against *Dmrtb1* and *Lrp8* in homozygous mutant and wild-type testis (*DMRTB1_B*^*-/-*^ *n*=2 and *DMRTB1_B*^*+/+*^ *n*=2). Both transcripts were expressed in different cell types throughout the cycle of the seminiferous tubules grouped into Early (I-V), Mid (VI-early VIII) and Late (late VIII-XII) spermatogenesis stages (Figures 2 and 3). In wild-type mice (*Dmrtb1_B*^*+/+*^), *Dmrtb1* signal is high in round spermatid (rSPD) cytoplasm in early stages, decreasing in intensity as the cycle progresses. At Mid cycle, rSPD expression is at the lowest, and isolated mRNA clusters in pachytene spermatocytes (SPCs) are visible. The late stages show strong signal in the cytoplasm of the second generation of SPCs (late Pachytene/Diplotene), but not the first generation (Leptotene/ Zygotene). The signal increases with progression of the cycle and strong signal can be seen in the nuclei of meiosis I SPCs. These observations indicate that *Dmrtb1* expression starts at pachytene SPCs, peaks at Diplotene and meiosis I SPCs, steadily decreases through rSPD and elongating spermatids (eSPDs) and is mostly absent at late eSPDs stages (Fig. 2A). The testis of *Dmrtb1_B*^*-/-*^ mice show a clear increase in *Dmrtb1* expression at all stages, without any apparent change in spatial and cell-type distribution (Fig. 2B). *Lrp8* also shows dynamic expression throughout the cycle of the seminiferous epithelium (Fig. 3A). Expression is higher in early stages and steadily reduces throughout the cycle, as a reflection of most signal being detected at the nuclei of rSPDs. Additionally, *Lrp8* foci can be detected in Sertoli cell (SC) nuclei, with some scattered signal in the surrounding cytoplasm. Similar to *Dmrtb1*, it appears that the main difference in *Lrp8* expression between genotypes is a slightly stronger overall signal in testes of mutant mice. The visual overexpression of both transcripts in testis of *Dmrtb1_B*^*-/-*^ mice was confirmed by image analysis of testis cross-sections (*n*=60 tubules/genotype/probe), which showed a small, but significant, increase in expression of *Dmrtb1* (p< 10^-4^; Wilcoxon test, Fig. 2B) and *Lrp8* (p< 10^-31^; Wilcoxon test; Fig. 3B).

**Figure 2.**
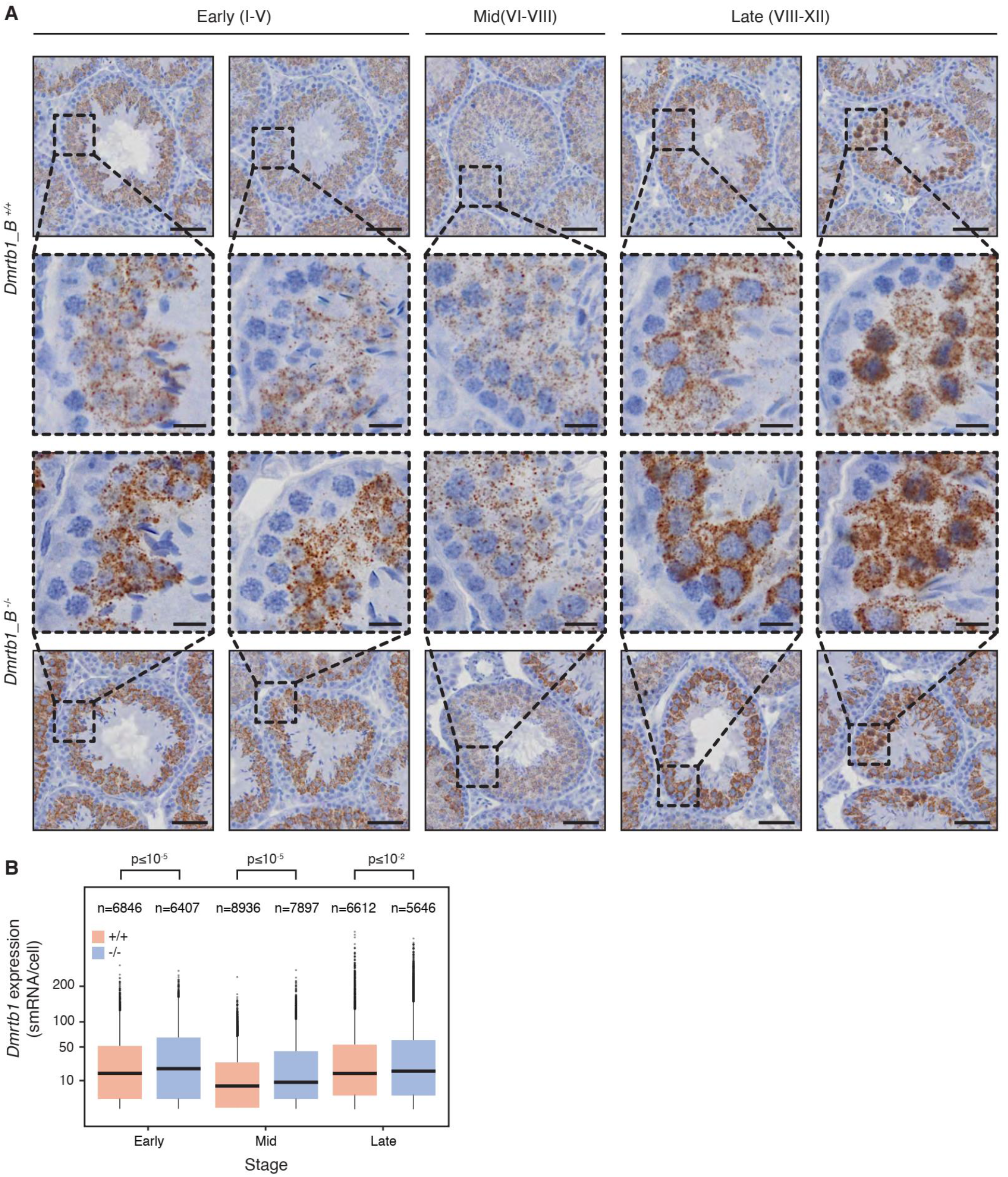
Expression of *Dmrtb1* in testis of wild-type and mutant mice. A) *Dmrtb1* mRNA expression was detected in testes sections of wild-type (*Dmrtb1_B*^*+/+*^, n=2) and mutant (*Dmrtb1_B*^*-/-*^, n=2) mice using RNA *in situ* hybridization. Examples of tubules in Early (I-V), Mid (VI-VIII), and Late (VIII-XII) stages are shown, as well as zoomed views of the tubule epithelium (red dash-lined boxes). Scale bar in full view = 50 µm, scale bar in zoomed view = 10 µm. B) *Dmrtb1* signal was quantified for a subset of tubules (n=120: 60 mutant + 60 WT) using the non-parametric Wilcoxon test to assess significance. The number above each box refers to the number of cells evaluated for the respective condition. smRNA: single-molecule RNA.

**Figure 3.**
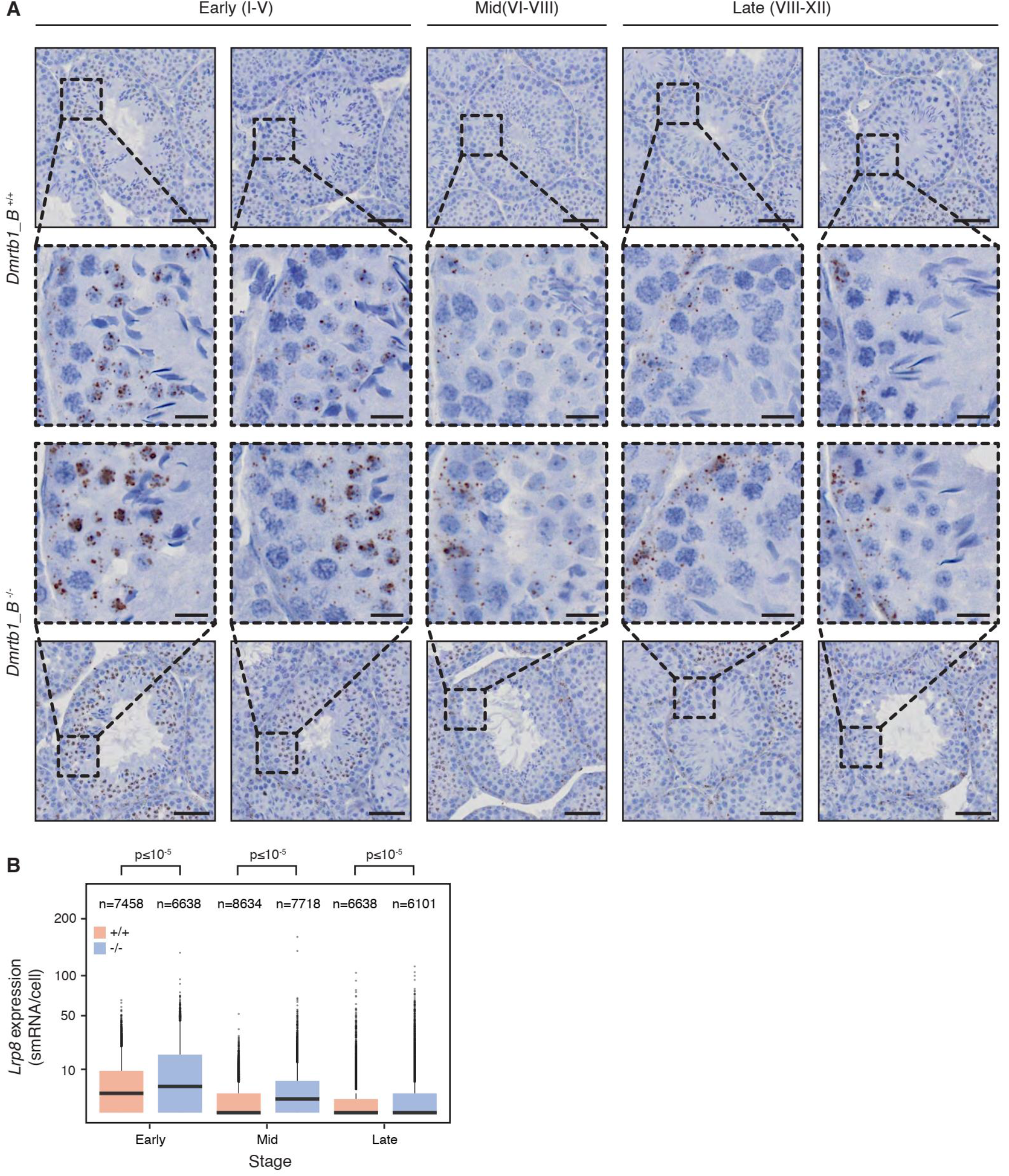
Expression of *Lrp8* in testis of wild-type and mutant mice. A) *Lrp8* mRNA expression was detected in testis sections of wild-type (*Dmrtb1_B*^*+/+*^, n=2) and mutant (*Dmrtb1_B*^*-/-*^, n=2) mice using RNA *in situ* hybridization. Examples of tubules in Early (I-V), Mid (VI-VIII), and Late (VIII-XII) stages are shown, as well as zoomed views of the tubule epithelium (red dash-lined boxes). Scale bar in full view = 50 µm, scale bar in zoomed view = 10 µm. B) *Lrp8* signal was quantified for a subset of tubules (n=120: 60 mutant + 60 WT) using the non-parametric Wilcoxon test to assess significance. The number above each box refers to the number of cells evaluated for the respective condition. smRNA: single-molecule RNA.

### Mutant mice have mild impairment of spermatogenesis

To investigate the potential impact of our TAD boundary deletion on fertility and viability, we examined litter size and genotype distribution among our heterozygous *Dmrtb1_B* crosses (Methods). We did not observe any effect on litter size (Student’s two-tailed t-test, p-value =0.225; Fig. 4A) or genotype distribution (*X*^*2*^ (2)= 0.316, p-value= 0.85; Fig. 4B), suggesting deletion of the target boundary has negligible impact on vitality and fertility, or that potential effects are only evident in other developmental, physiological or environmental contexts (e.g., advanced age, stress, poor diet, etc.).

**Figure 4.**
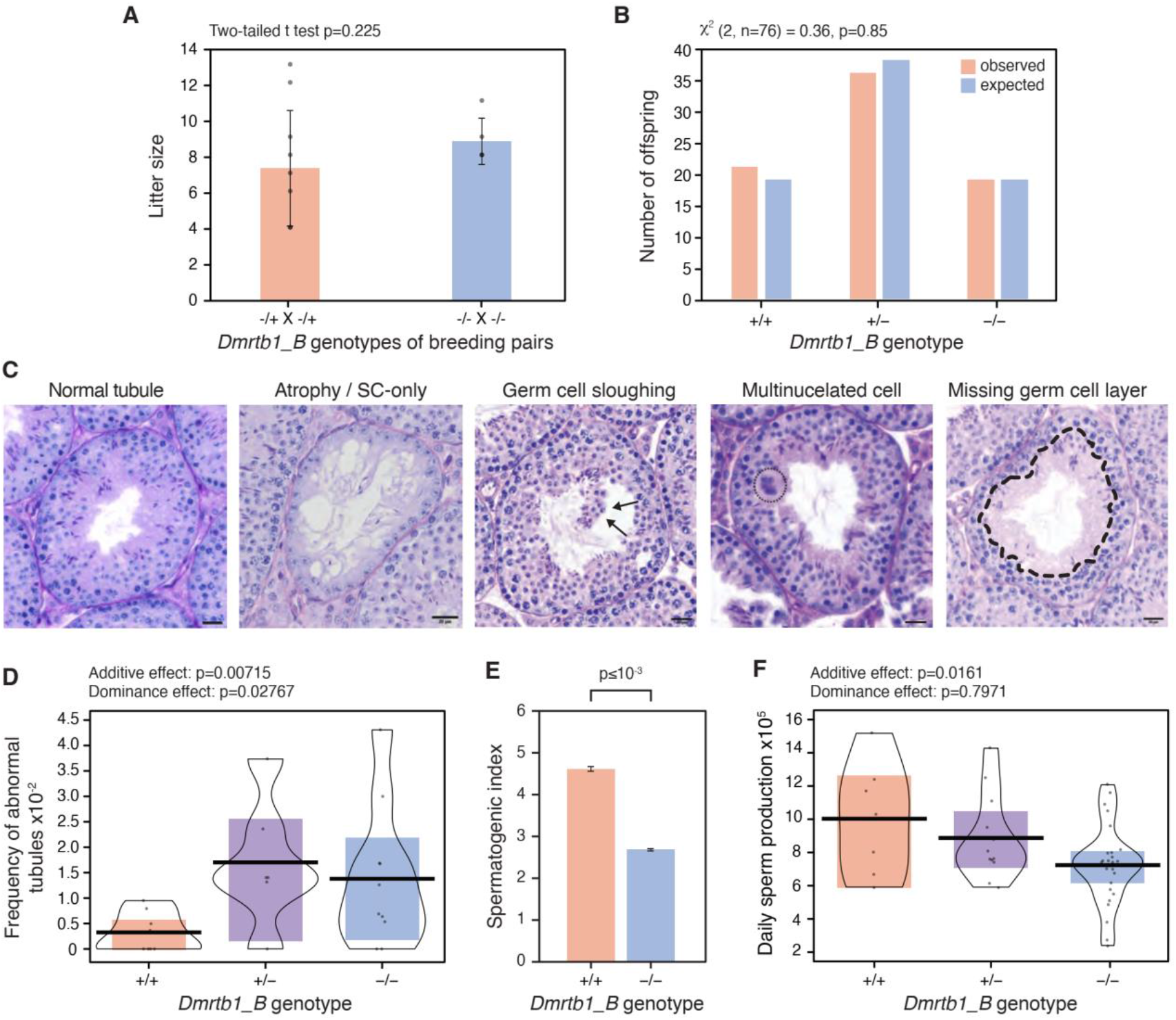
Evaluation of testicular histology and function of *Dmrtb1_B* mice. A) Litter sizes obtained from crosses of heterozygous (-/+, n=10) and homozygous (-/-, n=5) *Dmrtb1_B* mice. B) The observed and expected genotype distribution of offspring from heterozygous *Dmrtb1_B* cross. C) Example of a normal tubule along with different abnormal tubule phenotypes classified as severe in histopathology evaluation. D) Frequency of severe abnormalities in testis tubules from different genotypes (n=5,248). E) The spermatogenic index, defined as the ratio of round spermatids to spermatogonia, is shown as an estimate of testicular function (n= 3,464 tubules). A smaller index indicates more germ cell loss during the development of spermatogonia into spermatids. F) Daily sperm production estimates are shown (n=38).

Next, we asked whether the deletion of the conserved boundary is associated with changes in testis function by evaluating testis histology and daily sperm production (Fig. 4C-F). A total of 5,248 tubules (1,891 in *Dmrtb1_B*^*-/-*^, 1,304 in *Dmrtb1_B*^*+/-*^ and 2,053 in *Dmrtb1_B*^*+/+*^) were manually annotated for abnormality (Methods). Briefly, each tubule was annotated as either normal or abnormal (“spaces”, “vacuoles”, “apoptosis”, and “severe defects”; Supplementary Table 2). The severe defect category comprised tubules exhibiting the classical phenotypes of “Atrophy”, “Germ cell sloughing”, “Multinucleated cells” or “Missing germ cell layer” (examples shown in Fig. 4C). Evaluation of Periodic Acid-Schiff (PAS) stained tubules revealed a statistically significant increase in the frequency of severely abnormal tubules in testis of heterozygous and homozygous mutant mice, compared to wild-type (p= 0.007; Wald test), with evidence for dose dependent effect of the mutated allele (p= 0.028; Wald test) (Fig. 4D). Furthermore, automated image analysis of immunofluorescent testis cross-sections (two testis sections per *Dmrtb1_B*^*-/-*^ (*n*=3) and *Dmrtb1_B*^*+/+*^ (*n*=5) animals; total of *n*= 3,464 tubules) using SATINN^23^ (Fig. 4E) showed a 35% decrease in efficiency of spermatogenesis in homozygous mice (4.05 ± 0.048 to 2.63 ± 0.038, mean ± SE; p< 10^-5^), matching the 28% decrease estimated by daily sperm production (Fig. 4F; p=0.016; Wald test) from frozen testis of the same mice. Taken together, these data show subtle, yet significant, quantitative impairment of sperm production in mice homozygous for the *Dmrtb1_B* deletion.

## DISCUSSION

The mechanisms contributing to impaired spermatogenesis and male infertility have long been the subject of intensive investigation. However, the role of genome conformation, particularly Topologically Associated Domains (TADs), which play a pivotal role in regulating gene expression, remains largely unexplored. To examine the importance of TAD organization in the context of spermatogenesis, in this study we used genome editing to disrupt a TAD boundary *in vivo* and investigate its consequences at the molecular, histological, and functional level in the testis of a mouse model.

Since evolutionary conserved genomic elements are more likely to have an important functional role, our TAD boundary selection was guided by cross-species conservation of chromatin conformation. The conserved TAD boundary we selected for deletion was located near *Dmrtb1* and *Lrp8*, two genes implicated in spermatogenesis, suggesting it contributes to regulating sperm production processes. While several complementary assays confirmed successful deletion of our targeted DNA and the corresponding CTCF binding site, the deletion only resulted in a mild increase in DNA interaction frequency between the two adjacent TADs in testis of mutant mice. Other studies have also reported minimal disturbance in the local TAD organization despite deleting a TAD boundary or CTCF binding sites, indicating that additional mechanisms, apart from CTCF binding, contribute to maintaining TAD boundaries^24,25^. Alternatively, our use of bulk testis tissue may have obscured our ability to detect post-deletion changes in genome conformation, especially if TAD organization around our deletion region varies significantly across cell types or spermatogenesis stages, as reported previously^16,26^. Lastly, lack of drastic change in genome conformation following our deletion could indicate that functionally redundant CTCF binding sites are present nearby to provide functional stability and resilience in face of disruptions. Consistently, there are several CTCF sites located within 80 kb of the one we targeted, with two sites being located just 12kb upstream (Supplementary Fig. 1). In line with this, comparative studies have reported clustering of–possibly redundant– CTCF binding sites at and nearby TAD boundaries^27^, as well as a positive correlation between evolutionary conservation of TAD boundaries versus its CTCF binding sites and insulation strength^13^.

Despite the mild effect on chromatin interactions, both RNA-seq data (Fig. 1D) and quantification of mRNA *in situ* hybridization (ISH; Figs. 2B and 3B) showed overexpression of *Lrp8* and *Dmrtb1* transcripts in testis of our mutant mice. These data were statistically significant, except for *Lrp8* RNA-seq measurements, which was possibly too low to be detected in bulk tissue, as evidenced by the significant increase detected in the ISH data. Since these genes are known to play a role in spermatogenesis, we considered that even their mild overexpression could impact sperm development. Indeed, previous studies report *Dmrtb1* knock-out mice to be infertile and exhibit testicular maturation arrest, with failure of spermatogonia to enter meiosis in BL6 background mice, and spermatocytes failing to exit meiosis in 129v background^19^. In our study, *Dmrtb1* transcripts in wild-type mice were mostly expressed in late spermatocytes and round spermatids (Fig. 2A), consistent with mouse gene expression data^22,28^. Our ISH probes were designed to capture all *Dmrtb1* transcript isoforms, irrespective of coding potential, and should thus, reveal all cell-types expressing this gene during spermatogenesis. Importantly, we found that while maintaining cell-type specificity, mice lacking the TAD boundary have a minor increase in *Dmrtb1* expression throughout the cycle of the tubules (Fig. 2B). Given that DMRTB1 is a transcription factor capable of binding over 10K genes^19^, it is possible that even a mild overexpression could disrupt downstream processes, as seen in other overexpression phenotypes^29^. LRP8, also known as apolipoprotein E receptor 2 (APOER2), is the testis receptor for selenium-rich plasma protein selenoprotein P (SEPP1) and a component of the selenium delivery pathway to spermatogenic cells. Olson and colleagues have shown that mice lacking LRP8 have a sharp reduction in selenium levels and show sperm defects similar to those of *Sepp1*^*-/-*^ mice^21^. While the authors suggest that *Lrp8* expression is restricted to SCs, we found clear signal of *Lrp8* transcripts in both SC cytoplasm and rSPD nuclei (Fig. 3A), possibly from capturing novel transcript isoforms. Considering the role of this gene in selenium regulation and its importance for proper sperm morphology, even mild overexpression of *Lrp8* in *Dmrtb1_B*^*-/-*^ may lead to an increase in selenium uptake by spermatogenic cells, interfering with spermatid development and potentially leading to the reduced sperm counts we observed.

Our mutant *Dmrtb1_B*^*-/-*^ mice also showed a small increase in frequency of tubules with severe abnormalities, and a 28-35% decrease in daily sperm count, validated by two different methods (Fig. 4D-F), indicating that overexpression of *Dmrtb1* and *Lrp8* due to our deletion interferes with regulation of spermatogenesis. However, it is not clear whether the observed phenotypes result from interfering with expression of one of these genes, or both. Moreover, despite the disruption to spermatogenesis, we did not detect any evidence for reduced reproductive success in our mutant lines in the lab, such as failure to reproduce, skewed genotype ratios or birthing smaller litters. This suggests that the impact of our boundary deletion on fertility is negligible or is mitigated by compensatory mechanisms. It is also possible that reduced fertility is present but only detectible under unfavorable environmental or physiological conditions, such as poor lifestyle, poor health, or a more deleterious genetic background. Future comprehensive assays that use single-cell technology and investigate fertility of mutants across various contexts (e.g., advanced age, deletion of multiple CTCF binding sites, or environmental exposures) would help further reveal potential implications of TAD disruption for male reproductive health and infertility. Nonetheless, our work highlights the importance of considering non-coding functional variation, particularly in the context of TAD organization, as it can have subtle but significant effects on spermatogenesis. As demonstrated by this study, analysis of TAD organization in the context of spermatogenesis not only enhances our understanding of reproductive mechanisms but may also hold promise for making advancements in diagnosis and treatment of infertility in the future.

## METHODS

### Generating and validating Dmrtb1_B^-/-^ knockout mice

Our candidate TAD boundary was selected from a set of 18 boundaries previously reported^12^ to overlap synteny breakpoints between human and the *Nomascus leucogenys* gibbon, and to be evolutionarily conserved based on Hi-C data from the following species: human (lymphoblastoid cell lines or LCL), gibbon (LCL), rhesus (liver), rabbit (liver), dog (liver) and mouse (liver). The coordinates of the deletion were selected to include a CTCF binding site (chr4:107,737,644-107,738,425; mm10) that is conserved across mouse, rhesus, and human (Fig. 1A). Mice harboring the deletions (*Dmrtb1_B*^*-/-*^) were generated at the Transgenics Mouse Models core at OHSU by targeting a 23kb region (chr4: 107,716,841-107,739,906) by using six guide RNAs (gRNAs; left: L1, L2, L3; right R1, R2, R3; Supplementary Table 1, Supplementary Fig. 2) designed and tested in a genomic cleavage detection (GCD) assay in Neuro 2A murine cells by Thermo Fisher (Carlsbad, CA). Based on the results of the validation, L1, L3, R1, and R3 were selected for the boundary deletion. Knockout mice were produced using all four sgRNAs simultaneously and via the electroporation of one-cell-stage C57BL6/NJ JAX® mouse embryos using a NEPA 21 electroporator (NEPA GENE Co. Ltd., Chiba, Japan) as previously described^30^. The cells were also transfected with mRosa26 IVT gRNA as a positive control, and the TrueCut Cas9 protein v2 only, as negative control. Ribonucleoprotein complexes of SpCas9 protein (New England Biolabs) and gRNAs were prepared with a final concentration of 200 ng/μl and 25 ng/μl for each gRNA (100 ng/μl in total), respectively. After electroporation, embryos were transplanted into pseudo-pregnant recipient CD1/NCrl (Charles River) female mice.

To validate deletions in surviving offspring, genomic DNA was extracted from mouse tail clippings and PCR amplified using the KAPA Mouse Genotyping Kit (Roche) following manufacturer’s protocol. PCR amplicons were visualized on agarose gel to determine genotype. The primers and expected band sizes can be found in Supplementary Table 1. Breakpoints of the deletions were also confirmed using Sanger sequencing and Southern blot (Supplementary Fig. 2). A single mouse line with confirmed deletion was selected to generate all three genotypes: *Dmrtb1_B*^*-/-*^, *Dmrtb1_B*^-/+^, *Dmrtb1_B*^+/+^

### Generation and analysis of Capture Hi-C data

We used bulk testis from adult *Dmrtb1_B*^*-/-*^ and *Dmrtb1_B*^+/+^ (*n*=3 per genotype) to generate Capture Hi-C data. Hi-C libraries were generated using the Arima Hi-C Kit (Arima), following manufacturer’s protocol. Briefly, frozen fixed testes were lysed and conditioned before chromatin digestion. The digested chromatin was then filled in and biotinylated before ligation. Next, chromatin was protein-digested and reverse-crosslinking overnight, followed by purification. The purified DNA was then sonicated using the bioruptor pico (Diagenode) and size selected before library preparation using the NEB DNA Ultra II (New England Biolabs), following Arima’s protocol. The Hi-C DNA was bound to streptavidin beads before enzymatic end prep, adaptor ligation, DNA release by heat incubation, and lastly, PCR to barcode and amplify the libraries. Capture Hi-C libraries were generated by using the SureSelect Target Enrichment Kit (Agilent) and hybridizing the Hi-C libraries with custom probes before enrichment via Dynabeads MyOne Streptavidin Beads T1 (ThermoFisher). After library enrichment, a post capture PCR was conducted using 14 cycles, and libraries were paired-end sequenced at OHSU Massively Parallel Sequencing Shared Resource (MPSSR). Raw sequencing data was processed as described before^13^. Briefly, capture Hi-C data was aligned to *mm10* genome using HiCUP^31^. Pairtools (https://github.com/mirnylab/pairtools) was used to further process data and merge interaction matrices across replicates in each genotype. We then used HiCExplorer^32^ to obtain insulation scores along the targeted loci and compare interaction matrices between KO and control.

### Generation and analysis of CTCF chromatin immunoprecipitation sequencing (ChIP-seq) data

We used testis from adult *Dmrtb1_B*^*-/-*^ and *Dmrtb1_B*^+/+^ (three replicates per genotype) to generate CTCF ChIP-seq data. 50-100 mg of lightly chopped testis tissue was fixed with formaldehyde, quenched with glycine and then needle homogenized. The tissue slurry was pelleted and then lysed with 100ul of Lysis Buffer per 10 mg of tissue. Lysates then followed ChIP-seq protocols, as previously described^12^. We used 10 ul CTCF antibody (3418s; Cell Signaling Technologies). ChIP-seq libraries were prepared using the NEBNext Ultra II DNA Library Prep Kit for Illumina (New England Biolabs) without size selection, and libraries were sequenced on the Illumina HiSeq2500 or NovaSeq6000. After QC (Quality Control), reads were aligned to their respective genome assemblies using bowtie2^33^, with default settings. Reads with mapping quality <30 were removed. We also used MACS2^34^ (--nomodel --extsize 300) to generate tracks of log2 fold-enrichment (against input) for CTCF ChIP-seq data.

### Generation and analysis of RNA sequencing (RNA-seq) data

To examine changes in gene expression in *Dmrtb1_B*^*-/-*^ mice, we generated RNA-seq data from the testis. Briefly, after extracting total RNA from the testis (three replicates per genotypes) using the Monarch total RNA mini prep Kit (New England Biolabs), we generated RNA-seq libraries using the NEBNext Rrna Depletion Kit (New England Biolabs) and NEBNext Ultra II Directional RNA Library Prep Kit for Illumina kit (New England Biolabs), according to manufacturer’s instructions. Libraries were sequenced at MPSSR and reads were aligned to the mm10 genome using STAR^35^. The raw gene counts table generated and normalized to Transcript Per Million (TPM) values using custom R scripts. Routine differential analysis using DEseq2^36^ did not identify any significant changes in gene expression (p<0.05). However, since the TAD boundary deletion is expected to mostly influence expression of nearby genes and a function of the deletion genotype, we also used an eQTL approach to separately examine changes in the expression of genes nearby and distal to the deletion. Individuals homozygous for the deletion were assigned genotype 2, those heterozygous for deletion were considered 1 and wild-type individuals were 0. We used the linear regression method from MatrixEQTL^37^ and considered genes >100 kb from the center of the deletion to be in “cis” with the genotype locus (i.e., deletion). All other genes were considered is “trans”. Results with p<0.05 and FDR<0.05 were considered statistically significant for *cis* and *trans* linear regressions, respectively.

### Testis histological assays

To investigate testis histology and sperm development, freshly dissected gonads were collected from 8-week-old *Dmrtb1_B*^-/-^, *Dmrtb1_B*^-/+^, and *Dmrtb1_B*^+/+^ (i.e., wild-type) mice, and fixed under agitation in modified Davidson’s fixative (Electron Microscopy Sciences 64133-50) for 24 hours. Fixed tissues were washed in 70% Ethanol, embedded in paraffin, and sectioned at 5 μm. To capture images, we used an Olympus VS120 bright field, and fluorescent slide scanner equipped with DAPI, EGFP, Cy3, Cy5, and Cy7 filters (BrightLine® Sedat filter set DA/FI/TR/Cy5/Cy7-5X5M-B-000) with a 40X (numerical aperture 0.95; 0.17 mm/pixel) objective.

### Periodic Acid-Schiff stain (PAS)

Testicular cross-sections were stained with the Periodic Acid-Schiff kit (Epredia™; 87007) following the manufacturer’s standard staining protocol. Full sections of 24 mice (*n*=8 *Dmrtb1_B*^*-/-*^, *n*=6 *Dmrtb1_B*^*+/-*^ and *n*=10 *Dmrtb1_B*^*+/+*^) were scanned and analyzed blindly. A total of 5,248 tubules (1,891 in *Dmrtb1_B*^*-/-*^, 1,304 in *Dmrtb1_B*^*+/-*^ and 2,053 in *Dmrtb1_B*^*+/+*^) were evaluated and annotated as either normal or abnormal. Abnormal tubules were categorized as: 1) “apoptosis” (three or more apoptotic cells), 2) “spaces” (spaces in the tubule epithelium as a result of Sertoli cells detaching from one another), 3) “vacuoles” (one large vacuole, the size of the spermatocyte, or multiple smaller vacuoles within the epithelium) or 4) “severe defects” (multinucleated cells, atrophy (Sertoli cell only), missing germ cell layers, or sloughing of germ cells in the center of the tubule, Fig. 4C). Statistical analysis was performed in R v.4.2.0, using Poisson regression to compare the proportion of abnormal to normal tubules between genotypes. To account for sample batch effects, these comparisons were made fitting a generalized linear mixed model with genotypes as fixed and sample ID as random effects. When including all classes of tubule abnormalities (Supplementary Table 2), no statistically significant differences were found between genotypes. Therefore, we focused only on tubules with “severe defects” for the analysis represented in figure 4.

### Immunofluorescence

To perform automated image analysis of testis sections, we obtained immunofluorescence images on two testis sections per *DMRTB1_B*^*-/-*^ (*n*=3) and *DMRTB1_B*^*+/+*^ (*n*=5) animals. Specific protein markers were used to aid in boundary detection (anti-ACTA2, 1:100, Santa Cruz, sc-32251) and classification (anti-ACRV1, 1:200, Proteintech, 14040-1-AP) of tubules. Sections were deparaffinized in Xylenes and rehydrated through a series of decreasing concentrations of Ethanol. Tissue was permeabilized in 0.25% Triton X-100 (in 1X PBS), heat-induced antigen retrieval was performed in Universal Antigen Retrieval Reagent (Abcam, ab208572) and incubated in blocking solution (10% normal donkey serum, 1% BSA, 1X PBS). Primary antibodies were incubated overnight at 4°C, secondary antibodies (1:500, Invitrogen, A21202 and A21207) for 1 hour at room temperature and tissue was counterstained with 1 µg/mL Hoechst 33342. Whole-testis cross-sections were analyzed with the Software for Analysis of Testis Images with Neural Networks (SATINN^23^) for a total of 3,464 tubules (n=1,164 *B396_BOS*^*-/-*^ and n=2,300 *B396_BOS*^*+/+*^).

### mRNA in situ Hybridization (ISH)

The RNAscope 2.5 assay brown (ACD; #322300) was used to detect *Dmrtb1* and *Lrp8* transcripts in testicular tissue. For both transcripts, two adjacent 5 µm paraffin sections were used per slide, one section with either a *Dmrtb1* probe (ACD 485741) or a *Lrp8* probe (ACD 416801) and one with no probe as a control. *UBC* (ACD 310771), *Polr2a* (ACD 312471), and *DapB* (ACD 310043) probes were used in an additional control slide as a positive high expression control, a positive moderate expression control, and a negative control, respectively. All slides had 1 hour of baking time and 15 minutes of both protease plus treatment and target antigen retrieval. All other steps followed the manufacturer’s guidelines.

Whole-testis sections were analyzed using QuPath v.0.3.2 ^38^ to quantify and compare *Dmrtb1* and *Lrp8* expression in homozygous mutant (n=2 *Dmrtb1_B*^*-/-*^) and wild-type (*n*= 2 *Dmrtb1_B*^*+/+*^). For each probe, a total of 120 tubules (60 mutant + 60 WT) were manually staged (10 per stage category per animal) using the following criteria for assignment: a) Early (I-V): 2 generations of spermatids, no pre-Leptotene SPCs; b) Mid (VI-early VIII): 2 generations of spermatids, eSPD lined at the lumen and type-B spermatogonia or pre-Leptotene SPC present; c) Late (Late VIII-XII): no rSPD present, 2 generations of SPCs. Cell and single molecule segmentations were performed following ACD’s guidelines for image analysis with QuPath (Technical note MK 51-154/Rev A/Date 12/21/2020). Color deconvolution vectors were defined for Hematoxylin (0.75389, 0.6479, 0.10898), *Dmrtb1* and *Lrp8* H-DAB signal (0.35222, 0.60338, 0.71545), and background (218, 218, 202). Cell segmentation was then performed on the Hematoxylin channel (background radius = 14 µm; threshold = 0.06; cell expansion = 5 µm) and subcellular particle detection was performed on the original image (probe detection = 0.12; spot size = 0.3 µm). This resulted in a table containing information that captured the number of single mRNA dots per cell for each tubule. Data and statistical analysis were then performed in R v.4.2.0 by comparing the number of dots per cell per stage between genotypes, using the non-parametric Wilcoxon test to assess significance^39^.

### Measurement of daily sperm production

Daily sperm production (DSP) of 38 mice (*n*=7 *DMRTB1_B*^*-/-*^, *n*=12 *DMRTB1_B*^*+/-*^and *n*=19 *DMRTB1_B*^*+/+*^) was quantified and compared. Decapsulated testis were weighed, minced, and transferred to a pre-wet 35 µm medicon (BD Biosciences) with 1 mL of DSP buffer (0.9% NaCl, 0.01% Sodium Azide, 0.05% Triton X-100 in Milli-Q H2O) was added to the medicon and run on the medimachine (Becton Dickinson) for 15 min. The suspension was filtered through a pre-wet 20µm filter (Sysmex) and spun down at 500xg for 5 min. The resulting supernatant was discarded, and the pellet was resuspended in 1 mL chilled DSP buffer. 0.4% Trypan blue at 1:1 ratio was added to counterstain spermatid heads, which were counted using a Neubauer chamber. The number of spermatids per testis were calculated by dividing the number spermatids in homogenate by sample weight (g) and multiplying by the full testis weight (g). Developing spermatids spend 4.84 days in steps 14-16 in the mouse, so the total number of spermatids were divided by 4.84 to calculate the daily sperm production. Statistical analysis was then performed in R v.4.2.0 to compare DSP values between the genotypes. Data was standardized (subtracted the mean and divided by standard deviation) to fit normality (confirmed by distribution of residuals) and Wald test was used to evaluate the effect of gene dosage (additive model) on DSP.

### Investigating effect of deletion on viability and fertility

To investigate lethality of the TAD boundary deletion, we genotyped all offspring obtained from heterozygous crosses (i.e., *Dmrtb1_B*^*+/-*^ X *Dmrtb1_B*^*+/-*^) using the PCR assay described above and compared the observed genotype ratios to Mendelian expected genotype ratios (1:2:1) using a two-tailed chi-square test with the GraphPad Chi-Square calculator (https://www.graphpad.com/quickcalcs/chisquared1.Chi-square/; accessed December 2023). To examine potential effects of the deletion on fertility, we compared litter sizes (at birth) between heterozygous (*Dmrtb1_B*^+/-^ X *Dmrtb1_B*^+/-^, n=10) and homozygous crosses (*Dmrtb1_B*^*-/-*^ X *Dmrtb1_*^*-/-*^, n=5) using a two-tailed t test.

## DATA ACCESSIBILITY

All testis CTCF ChIP-seq, RNA-seq and Capture Hi-C datasets generated as part of this study are available at the Gene Expression Omnibus (GEO), under the accession number GSE242235.

## ACKNOWLEDGEMENTS

The ChIP-seq assays were performed by the KCVI Epigenetics Consortium at OHSU. Next-generation libraries were sequenced at the OHSU Massively Parallel Sequencing Shared Resource and Novogene Co. Data analyses were performed on the Exacloud computer cluster at OHSU. This research was supported by the Integrated Pathology Core at the Oregon National Primate Research Center (ONPRC) which is supported by NIH Award P51 OD 011092. The content is solely the responsibility of the authors and does not necessarily represent the official views of the National Institutes of Health.

