## Supplementary material for "Deletion of an evolutionarily conserved TAD boundary compromises spermatogenesis in mice": Suplemental Data

Lima *et al.*

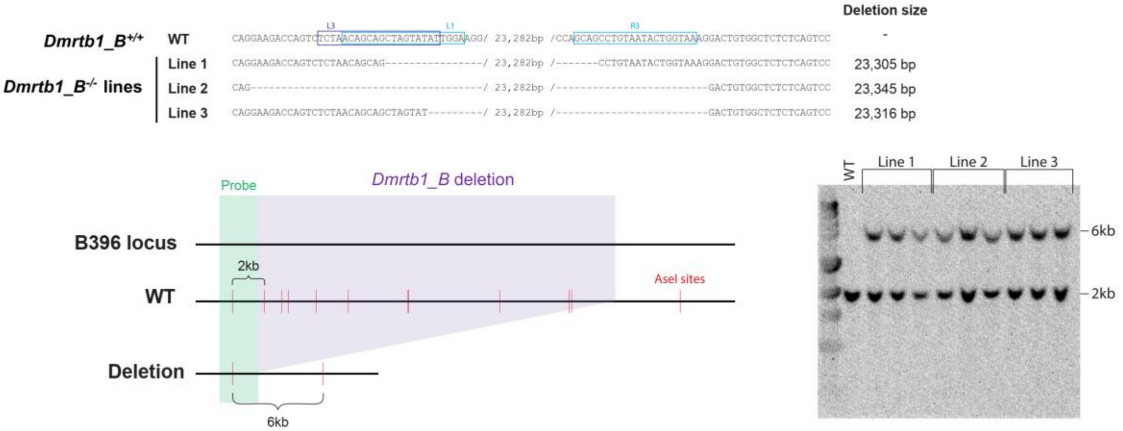


**Figure S1. Validation of the deletion in mutant mouse lines (*Dmrtb1_B*$^{-/-}$).** Top panel shows breakpoints of deletions per CRISPR mouse line, as determined with Sanger sequencing. Bottom panels show Southern blot validation of the deletions. Mouse line number 1 was selected for all downstream analysis.

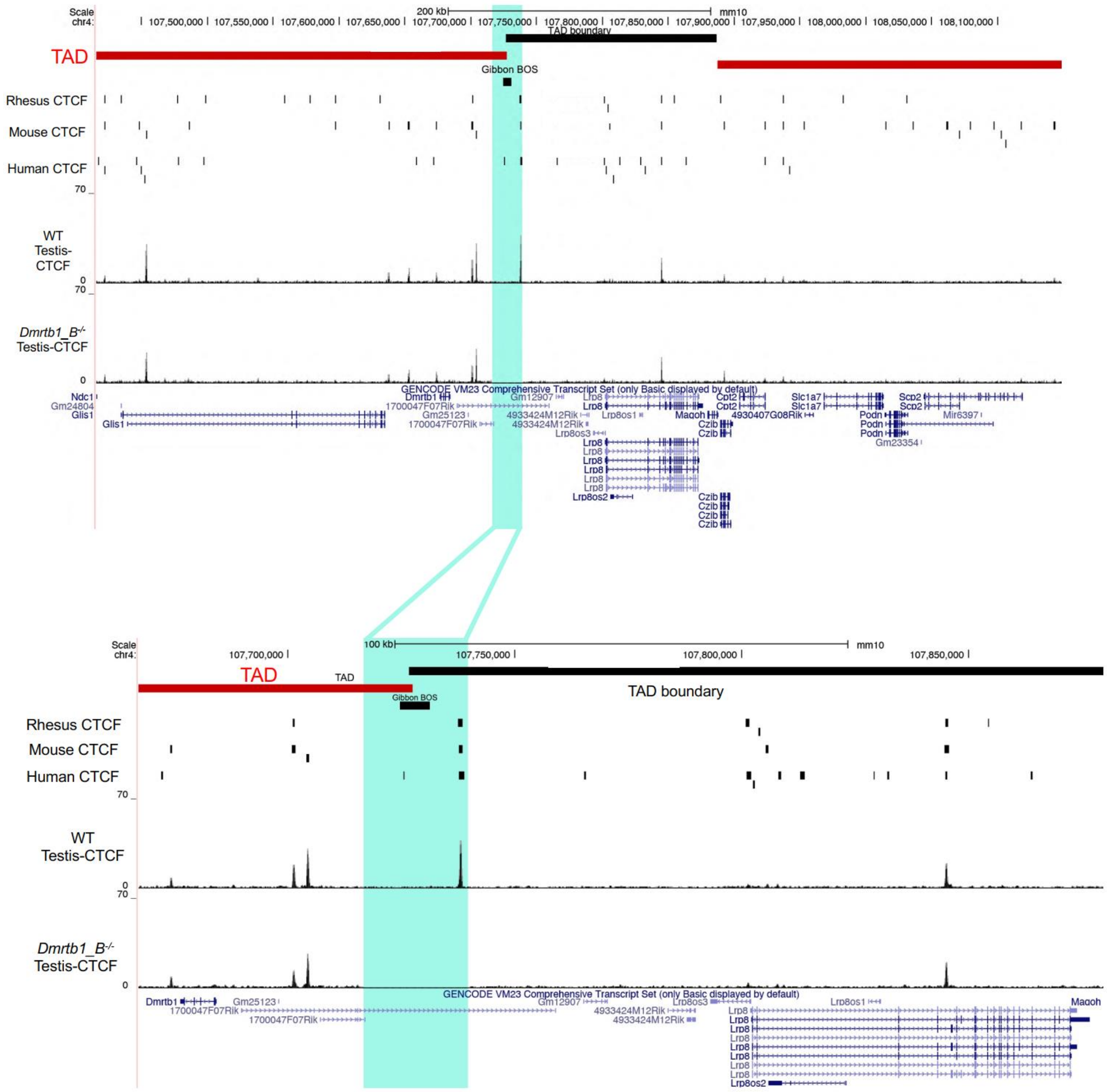


**Figure S2.** Upper panel shows a screenshot from the UCSC browser around the deleted region (shaded in aqua) within an evolutionarily conserved TAD boundary (black bar). The deletion includes a gibbon break of synteny (BOS) and an evolutionary conserved CTCF site (CTCF binding also evident in mouse testis based on our ChIP-seq data). Lower panel is a zoomed in image of the deletion site and demonstrates successful removal of CTCF binding in the mouse testis following of the CRISPR-Cas9 deletion.

**Table S1.** A list of all the guide RNAs (gRNA) and primers used to validate the deletion through Southern blot and Sanger sequencing.

| | **Guide RNAs** | | **Southern Blot** | | **Genotyping** | | | | | |
|---|---|---|---|---|---|---|---|---|---|---|
| **Line** | **gRNA names** | **gRNA sequences** | **Primer** | **Sequence (5'-3')** | **Primer** | **Sequence (5'-3')** | **Primer** | **Sequence (5'-3')** | **Expected band size w/ deletion** | **Expected band size w/out deletion** |
| ***Dmrtb1_B*** | L1 | ACAGCAGCTAGTATATTGGA | MmDel_B396_SB-F | CGAGGCTACCGTTCACCTTA | MmDel_B396_Gen-F1 | GAACTTGCCAAAGGACTGAG | MmDel_B396_Gen-R1 | TCCCTTAATCTCCATTTCCA | No band | 201 bp |
| | L2 | ATCAGACCAGCATTGTCTGC | MmDel_B396_SB-R | CATCACACAACCACCCTCAG | MmDel_B396_Gen-F2 | AGGAGGAGTCCAGAGTCCAG | MmDel_B396_Gen-R2 | GATTTGGATCAAGCTCCTGT | No band | 249 bp |
| | L3 | TCTAACAGCAGCTAGTATAT | | | MmDel_B396_Gen-F1 | GAACTTGCCAAAGGACTGAG | MmDel_B396_Gen-R2 | GATTTGGATCAAGCTCCTGT | 258 bp | No band |
| | R1 | TTACCAGTATTACAGGCTGC | | | | | | | | |
| | R2 | GTGACTAACAGTGACTCCTT | | | | | | | | |
| | R3 | GCAGCCTGTAATACTGGTAA | | | | | | | | |

**Table S2.** Classification and quantification of tubule abnormalities from histological images of PAS-stained testes of mice of different *Dmrtb1_B* deletion genotypes.

| Sample ID | Total # tubules | # abnormal tubules | Spaces | Vacuoles | Apoptosis | Severe defects | Genotype | Batch |
|---|---|---|---|---|---|---|---|---|
| 400M_Hom | 239 | 44 | 14 | 23 | 3 | 4 | HOM | 1 |
| 769M_Hom | 157 | 26 | 9 | 15 | 1 | 1 | HOM | 1 |
| 101M077 | 210 | 19 | 2 | 10 | 5 | 2 | WT | 1 |
| 86M078 | 200 | 40 | 28 | 5 | 1 | 6 | HOM | 1 |
| 85M078 | 225 | 9 | 5 | 3 | 1 | 0 | WT | 1 |
| Crispy | 175 | 33 | 16 | 16 | 1 | 0 | HOM | 1 |
| 83M078 | 196 | 23 | 12 | 9 | 2 | 0 | WT | 1 |
| 87M078 | 259 | 36 | 19 | 12 | 5 | 0 | WT | 1 |
| 128M756 | 214 | 16 | 3 | 4 | 1 | 8 | HET | 2 |
| 129M756 | 222 | 4 | 0 | 4 | 0 | 0 | HET | 2 |
| 130M756 | 252 | 48 | 15 | 32 | 1 | 2 | WT | 2 |
| 147M755 | 238 | 35 | 13 | 19 | 0 | 3 | HOM | 2 |
| 148M755 | 214 | 32 | 13 | 14 | 3 | 3 | HET | 2 |
| 149M755 | 243 | 51 | 11 | 44 | 1 | 0 | HOM | 2 |
| 150M756 | 145 | 3 | 1 | 1 | 0 | 1 | HOM | 2 |
| 157M006 | 214 | 26 | 11 | 12 | 0 | 3 | HET | 2 |
| 160M755 | 212 | 26 | 4 | 15 | 2 | 5 | HET | 2 |
| 161M755 | 228 | 11 | 5 | 3 | 0 | 3 | HET | 2 |
| 197M750 | 202 | 10 | 7 | 1 | 2 | 1 | WT | 2 |
| 203M755 | 237 | 23 | 9 | 10 | 0 | 4 | HOM | 2 |
| 204M755 | 232 | 33 | 8 | 15 | 0 | 10 | HOM | 2 |
| 211M766 | 272 | 17 | 7 | 10 | 0 | 0 | WT | 2 |
| 222M766 | 275 | 47 | 11 | 36 | 0 | 1 | WT | 2 |
| 100M071 | 187 | 32 | 9 | 20 | 2 | 1 | HOM | 1 |
